# Brain Structure and Episodic Learning Rate in Cognitively Healthy Ageing

**DOI:** 10.1101/2022.05.03.490431

**Authors:** Darya Frank, Marta Garo-Pascual, Pablo Alejandro Reyes Velasquez, Belén Frades, Noam Peled, Linda Zhang, Bryan A. Strange

**Affiliations:** Laboratory for Clinical Neuroscience, CTB, Universidad Politécnica de Madrid, 28223 Madrid, Spain; Alzheimer’s Disease Research Unit, CIEN Foundation, Queen Sofia Foundation Alzheimer Center, 28031 Madrid, Spain; PhD Program in Neuroscience, Autonoma de Madrid University, 28049 Madrid, Spain; Athinoula A. Martinos Center. for Biomedical Imaging, Department of Radiology, Massachusetts General Hospital, 02129 Charlestown, MA, USA; Harvard Medical School, 02115 Boston, MA, USA

## Abstract

Memory normally declines with ageing and these age-related cognitive changes are associated with changes in brain structure. Episodic memory retrieval has been widely studied during ageing, whereas learning has received less attention. Here we examined the neural correlates of episodic learning rate in ageing. Our study sample consisted of 982 cognitively healthy female and male older participants from the Vallecas Project cohort, without a clinical diagnosis of mild cognitive impairment or dementia. The learning rate across the three consecutive recall trials of the verbal memory task (Free and Cued Selective Reminding Test) recall trials was used as a predictor of grey matter (GM) using voxel-based morphometry, and WM microstructure using tract-based spatial statistics on fractional anisotropy (FA) and mean diffusivity (MD) measures. Immediate Recall improved by 1.4 items per trial on average, and this episodic learning rate was faster in women and negatively associated with age. Structurally, hippocampal and anterior thalamic GM volume correlated positively with learning rate. Learning also correlated with the integrity of WM microstructure (high FA and low MD) in an extensive network of tracts including bilateral anterior thalamic radiation, fornix, and long-range tracts. These results suggest that episodic learning rate is associated with key anatomical structures for memory functioning, motivating further exploration of the differential diagnostic properties between episodic learning rate and retrieval in ageing.

## Introduction

Ageing is accompanied by a decline in cognition, most characteristically in episodic memory performance (Glisky, 2007; Tromp et al., 2015), the ability to remember personal experiences. Episodic memory impairments in ageing can manifest in different ways depending on the studied phase (i.e., encoding, consolidation, or retrieval process). It is difficult to study these phases independently in behavioural studies, although previous work has reported that distinct processes may be affected unequally during ageing. For example, more prominent deficits have been found for encoding relative to retrieval in older adults (Friedman et al., 2007; Morcom et al., 2003). Furthermore, exploring the neural underpinnings of these various manifestations (e.g., learning versus retention) could inform dissociations between normal age-related decline and decline driven by neurodegenerative diseases such as dementia. Memory decline in ageing is often measured in terms of retention. However, impairments could also be driven by a diminished ability to learn information over a period of time, rather than to retrieve it. We therefore aimed to elucidate the structural brain properties underlying episodic learning rate in ageing.

Ageing has been associated with a global reduction in grey matter (GM) volume (Farokhian et al., 2017; Grieve et al., 2007), although to different extents across brain regions (Cox et al., 2018; Resnick et al., 2003). Numerous studies have found a specific GM volume loss in prefrontal, temporal and parietal cortices (Cox et al., 2021; Elliott, 2020), associated with general cognitive and memory-specific decline (Cox et al., 2021; Fjell and Walhovd, 2010; Gorbach et al., 2017). White matter (WM) age-related differences in fractional anisotropy (FA) and mean diffusivity (MD) have also been reported (Bennett et al., 2010; Fjell and Walhovd, 2010; Madden and Parks, 2017). FA and MD are negatively correlated such that reduced WM integrity is indexed by a decrease in FA and an increase in MD. Like GM, age-related WM effects are apparent throughout the brain (Farokhian et al., 2017; Grieve et al., 2007), although with greater effects in anterior than posterior tracts (Bennett et al., 2010). Whilst these structural differences contribute to our understanding of brain ageing, it is vital to also consider their cognitive manifestations.

A commonly observed form of age-related cognitive decline is impaired memory, which has been associated with reduced hippocampal volume (Gorbach et al., 2017; Hedden et al., 2016; Persson et al., 2012), as well as with damage to the microstructure of frontal and temporal WM tracts (de Mooij et al., 2018; Kennedy and Raz, 2009; Rizvi et al., 2020), and specifically limbic tracts (Bennett et al., 2015). Furthermore, recognition performance on neuropsychological episodic memory tests has been shown to correlate with FA and MD measures in the fornix, cingulum, and superior and inferior longitudinal fasciculi (Sasson et al., 2013). However, other studies have not found correlations between WM microstructure and episodic retrieval in ageing (Gorbach et al., 2017; Laukka et al., 2013; Salami et al., 2012).

Memory performance is usually quantified by the ability to recognise or recall information correctly, a retrieval impairment could be caused by a reduced ability to encode or learn information (Boujut and Clarys, 2016; Cadar et al., 2018). Encoding, which is potentially dissociable from retrieval processes (Bennett et al., 2015; Kwok and Buckley, 2010), has been shown to underlie several memory deficits observed in ageing (Grady, 2012). Whilst learning rate is part of the encoding process, in the current context it specifically refers to an improvement in learning over time (or repetitions). Indeed, there is evidence for reduced error-driven (Nassar et al., 2016) and probabilistic learning rates (Herff et al., 2019; Samanez-Larkin et al., 2012) in older adults, but evidence for similar deficits in episodic learning rate is lacking.

A potential way to probe episodic learning rate is through the Free and Cued Selective Reminding Test (FCSRT). The FCSRT is one of the most commonly used free-recall paradigms for episodic memory assessment, including immediate and delayed free- and cued-recall (Buschke, 1984). Worse recall performance of cognitively normal older adults on the FCSRT has been associated with reduced hippocampal GM volume (Zammit et al., 2017), reduced fornix FA (Hartopp et al., 2019; Metzler-Baddeley et al., 2011) and increased frontal MD (Nicolas et al., 2020). In addition to such retrieval effects, using the immediate free recall components across three consecutive trials, the FCSRT enables investigating episode learning by examining how many additional words are successfully recalled on each trial.

We investigated whether the learning rate in FCSRT is associated with age, as well as its neural manifestation in GM volume and WM tract microstructure. In a large cross-sectional cohort of healthy older adults, we first calculated the learning rate across the three consecutive FCSRT trials and tested for an association with age. To examine brain-cognition associations, we used learning rate as a predictor to examine 1) GM volume using voxel-based morphometry (VBM), and 2) WM microstructure using tract-based spatial statistics on FA and MD measures. Given the critical role of the hippocampus in episodic memory and its correlation with structural changes in ageing, we hypothesised that hippocampal GM volume and associated WM limbic tract microstructure would correlate with learning rate.

## Methods

### Participants

All participants in this study were part of the Vallecas Project, a single-centre longitudinal study of community-dwelling volunteers aged 69-86 without any cognitive or psychiatric disorder that compromised their daily functioning at the time of recruitment. Inclusion and exclusion criteria have been further described elsewhere (Olazarán et al., 2015). From this cohort, data from the baseline visit of 982 cognitively normal participants (mean age = 74.8, SD = 3.9, 637 (64.9%) females) were included in the current study. Any subject with a diagnosis of mild cognitive impairment or Alzheimer’s disease at this first visit was excluded. All participants provided written informed consent and the Vallecas Project was approved by the Ethics committee of the Instituto de Salud Carlos III.

### Neuropsychological assessment

Participants completed a battery of neuropsychological assessments as part of the Vallecas Project protocol. In this study, we report the total score of the Mini Mental State Examination (MMSE; Folstein et al., 1975) and we mainly focused on the Free and Cued Selective Reminding Test (FCSRT; Buschke, 1984), assessing learning and retention of verbal memory, with immediate and delayed recall components. The test was administered using standard procedures (Peña-Casanova et al., 2009). Participants were presented with cards containing four words and asked to identify the word corresponding to a specific semantic category, going through all four words, on four different cards (16 words in total). The words presented are not the most obvious member of each semantic category. Following the presentation phase, participants were asked to recall as many words as possible in three consecutive recall trials each one followed by 20 seconds of interference counting backwards (Figure 1A). For each trial, participants were asked to freely recall as many words as possible with a time limit of 90 seconds, then examiners provided the semantic category clue for the forgotten items. These three free and cued recalls constitute the three immediate recall trials of the task. This immediate recall phase is followed by a 30-minute delay, after which the delayed phase of the test starts. Participants were asked on a single trial to freely recall as many words as possible otherwise cues were provided (Figure 1A). To assess the learning rate across trials, we fit a linear mixed-effects model of the number of items freely recalled in each immediate trial, as a function of the recall trial (first, second, and third) using the lme4 package in R 4.0.2 (https://www.r-project.org/). The model also included a random slope of the recall trial, and a random intercept per participant, capturing inter-individual variability in learning rate (across the three trials). The learning rate coefficient for each participant was extracted using the coef() function for subsequent analyses. Next, we built a multiple regression model where the learning rate was the dependent variable, sex, age and level of education were the predictors and the delayed free recall score of the FCSRT was included as a covariate to rule out the retrieval phase of the memory process. Extraction and plotting of the effects reported below were conducted using the effects (Fox, 2003) and ggplot2 (Wickham, 2009) packages in R.

**Figure 1.**
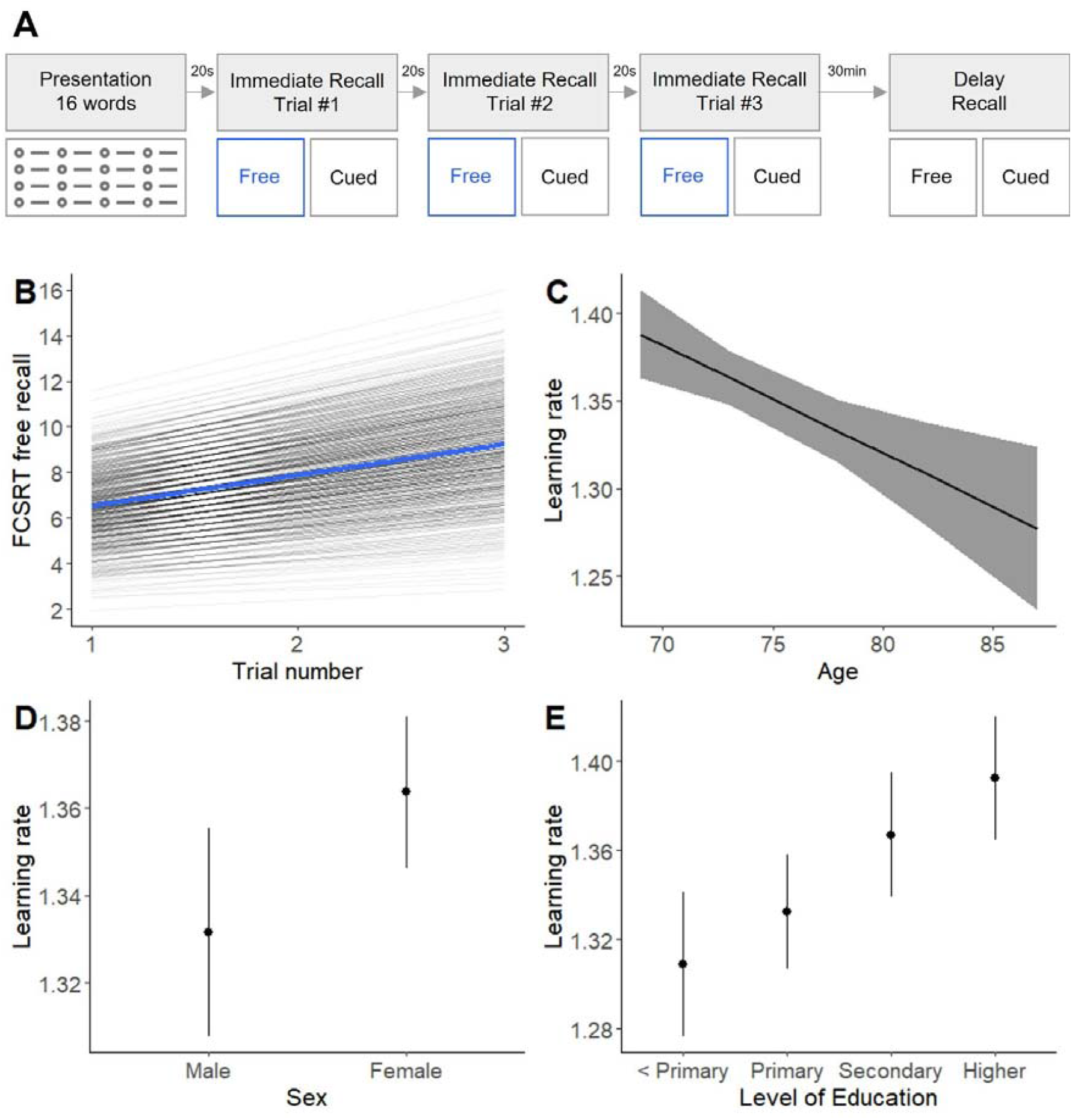
Learning rate across FCSRT trials is related to age, sex and level of education. A Diagram of the FCSRT protocol used to assess memory in this study in which the learning rate is calculated from the free recall of the three immediate trials (blue). B Mean and individual learning rates across the three free immediate recall FCSRT trials. After controlling for the rest of the model predictors, C learning rate decreases with age, D females learn faster than males and E level of education is positively associated with learning rate. Error bars represent 95% confidence intervals.

### MRI Data acquisition

Images were acquired using a 3T MRI (Signa HDxt GE) with a phased array eight-channel head coil. T1-weighted images (3D fast spoiled gradient echo with inversion recovery preparation) were collected using a repetition time (TR) of 10ms, echo time (TE) of 4.5ms, field of view (FOV) of 240mm and a matrix size of 288×288 with slice thickness of 1mm, yielding a voxel size of 0.5 × 0.5 × 1 mm3. Diffusion-weighted images were single-shot spin echo echo-planar imaging (SE-EPI), with TR 9200ms, TE 80ms, b-value 800s/mm2 and 21 gradient directions, FOV 240mm and matrix size 128 × 128 with slice thickness of 3mm.

### Grey matter VBM

The analysis was carried out in SPM12 (version r6225; https://www.fil.ion.ucl.ac.uk/spm). T1-weighted images were segmented into grey matter, white matter and cerebrospinal fluid and then aligned and normalised to MNI space using the DARTEL algorithm (Ashburner, 2007). Prior to statistical modelling, the normalised images were smoothed using a 6mm FWHM Gaussian kernel. The pre-processed grey matter maps were entered into a general linear model (GLM) with learning rate from the memory task as the predictor of interest, and total intracranial volume (TIV), sex, and the delayed free recall score of the FCSRT as covariates. Age and education were not used in the model as additional covariates since FCSRT delayed free recall is sensitive to the effects of age and level of education. Nonetheless, to ensure the model is capturing variance associated with these variables we devised a second model without FCSRT delayed free recall and including TIV, sex, age and education as covariates and the same results were obtained (see Supplementary Materials). We conducted whole-brain analyses using a threshold-free cluster enhancement (TFCE) approach with 5000 permutations and default parameters (E = 0.5 and H = 2) using the TFCE tool (version r223) for CAT12 toolbox in SPM (http://dbm.neuro.uni-jena.de/tfce). Therefore, our analyses fully correct for mass-univariate testing (and associated multiple-comparisons problem) by employing a whole-brain FWE correction. Furthermore, we used the TFCE approach to overcome cluster-based inference issues. The AAL3 atlas neuroanatomical labels were used to describe neuroanatomical loci (Rolls et al., 2020) and Mango software was used to produce the figure (http://rii.uthscsa.edu/mango/). These analyses assessed which regions were positively associated with the immediate recall learning rate. Significant results are reported at a family-wise error FWE) corrected level of p < 0.05.

### White matter tract-based spatial statistics (TBSS)

Of the 982 participants, seven were excluded as they did not have diffusion data. For preprocessing these images, the FSL toolbox (http://fsl.fmrib.ox.ac.uk/fsl/fslwiki) was used for motion and eddy current correction, the extraction of non-brain voxels and, lastly, the calculation of voxel-wise diffusion maps (FA and MD) for each participant. Individual FA and MD maps were then used in the FSL TBSS pipeline (http://fsl.fmrib.ox.ac.uk/fsl/fslwiki/TBSS/UserGuider; detailed methods described by Smith et al. (2006)). The general outline of the process is: 1) FA individual maps were non-linearly registered to standard space (FMRIB58_FA template) (Andersson et al., 2007). 2) A mean FA image was created by averaging all co-registered FA maps. 3) Individually aligned images were projected onto the mean FA skeleton—representing the centers of all tracts common to the study sample—and skeletonised images were used for voxel-wise analysis. Diffusivity maps for MD were generated by applying the same steps detailed above. The same GLM design matrix as the VBM analysis was used along with the TFCE approach with 5000 permutations (default parameters E = 0.5 and H = 2). Significant results are reported at a family-wise error (FWE) corrected level of p < 0.05. To visualise our TBSS results we used the multimodal analysis and visualisation tool (MMVT; Felsenstein et al., 2019). The pipeline follows these steps: 1) Binary masking: all the voxels in the TBSS volume below the threshold (0.95) were set to zero. 2) Outlier voxels removal using the Open3D python package (Zhou et al., 2018). 3) Smoothing the volumetric data using a 3D Gaussian filter (Virtanen et al., 2020). 4) Surface creation from the volume’s TBSS surfaces using the marching cubes algorithm (Lorensen and Cline, 1987). For that, we re-calculate the threshold to give us the same number of voxels after the smoothing step. 5) Translation for the surfaces’ vertices coordinates. 6) Projection of the volumetric data on the surfaces.

## Results

### Memory and neuropsychological performance

On average, across three trials, participants correctly remembered 7.9 items (SD = 2.6). When looking at individual trials, performance improved as trials progressed, reflecting a positive episodic learning rate (see Table 1 for number of items recalled, and Figure 1B for learning rates). Our linear model predicting the learning rate as a function of age, sex, and level of education revealed significant effects of the three predictors after correcting for FCSRT delayed free recall score. Learning rate and delayed free recall FCSRT were positively correlated (Pearson’s r = 0.7; p < 2.2×10-16). Age had a negative effect on learning rate (F(1,965) = 10.45, p = 0.001) (Figure 1C), sex also had an effect (F(1,965) = 4.66, p = 0.031) and being a woman was positively associated with learning rate (Figure 1D). Finally, having a higher level of education was positively associated with learning rate (F(3,965) = 5.17, p < 0.002) (Figure 1E). There was no significant interaction between the three predictors (age, sex and years of education).

**Table 1.**
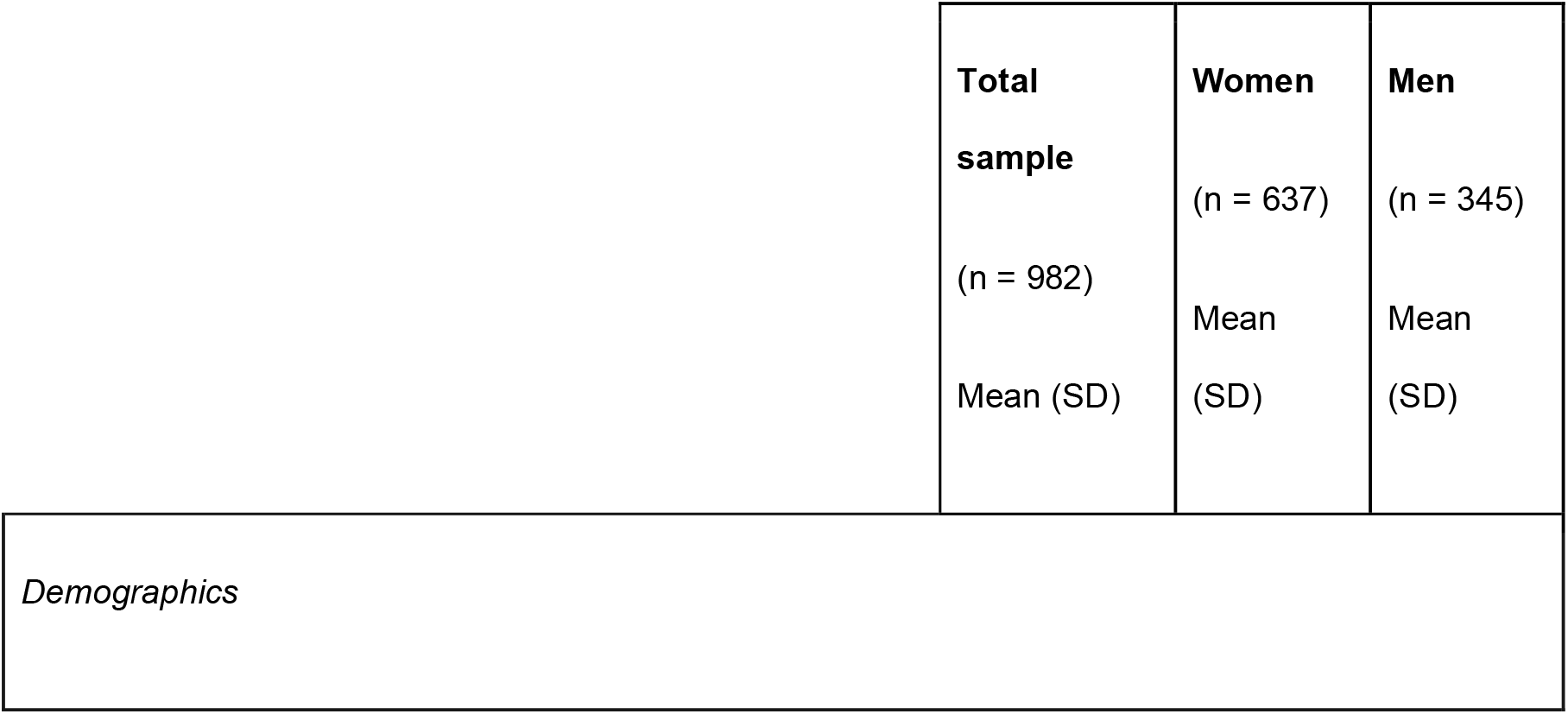
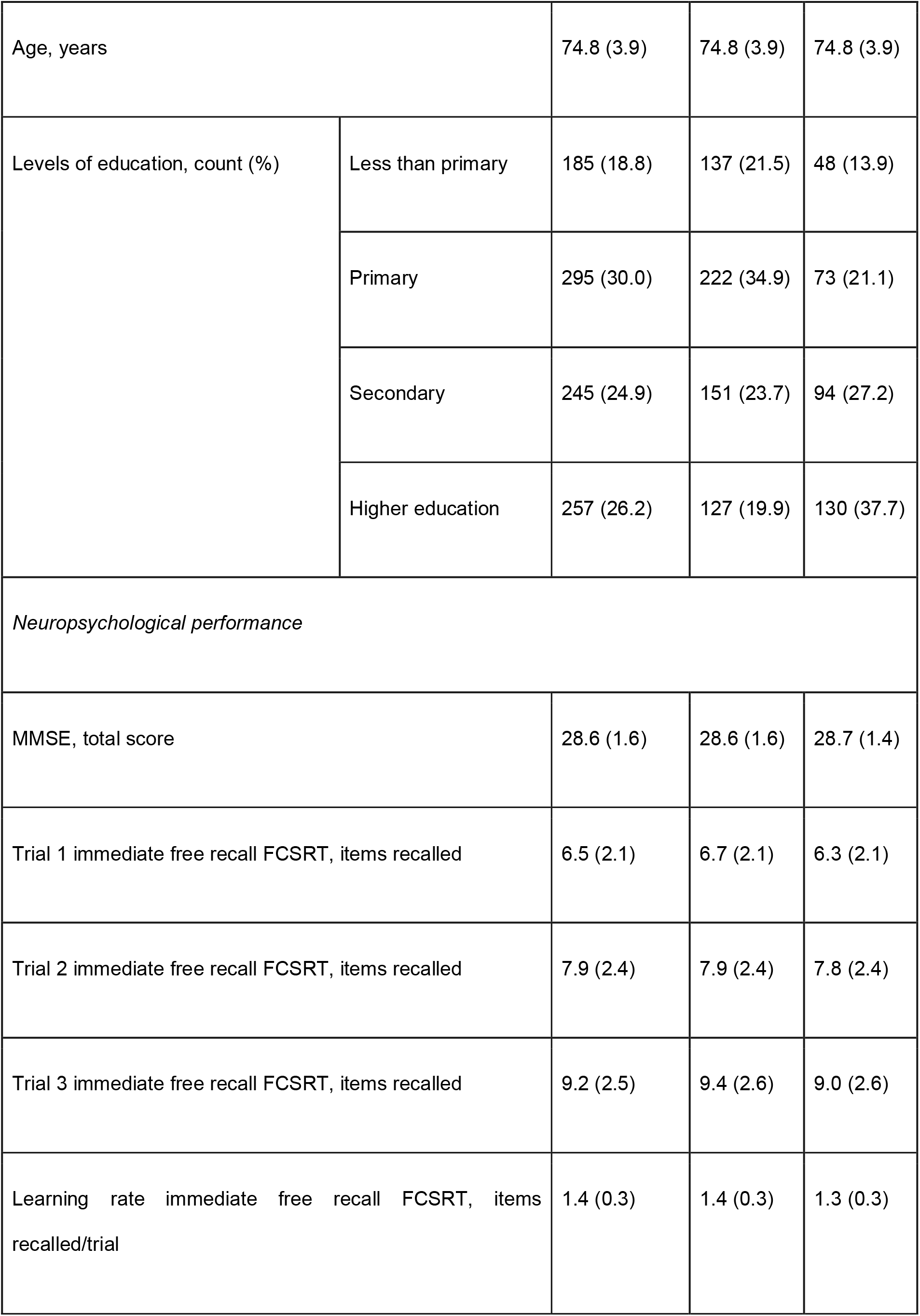
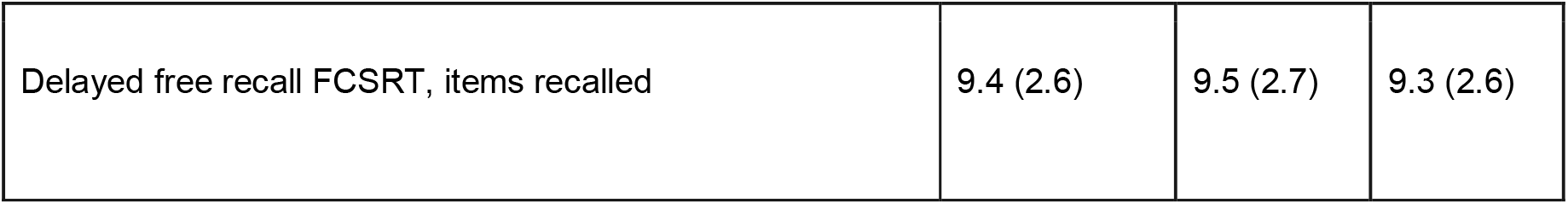
Demographic and neuropsychological profile of the total sample and split by sex. MMSE: Mini Mental State Examination total score, FCSRT: Free and Cued Selective Reminding Test.

### Grey matter volume (VBM)

We found a positive correlation between episodic learning rate and grey matter volume in the bilateral hippocampus, with more pronounced effects on the left side and the left superior temporal gyrus (Figure 2A, Supplementary Figure 1), and the right anterior thalamic nucleus with some extension to adjacent nuclei (right ventroanterior and ventrolateral thalamic nuclei; Figure 2B, Supplementary Table 1, Supplementary Figure 1).

**Figure 2.**
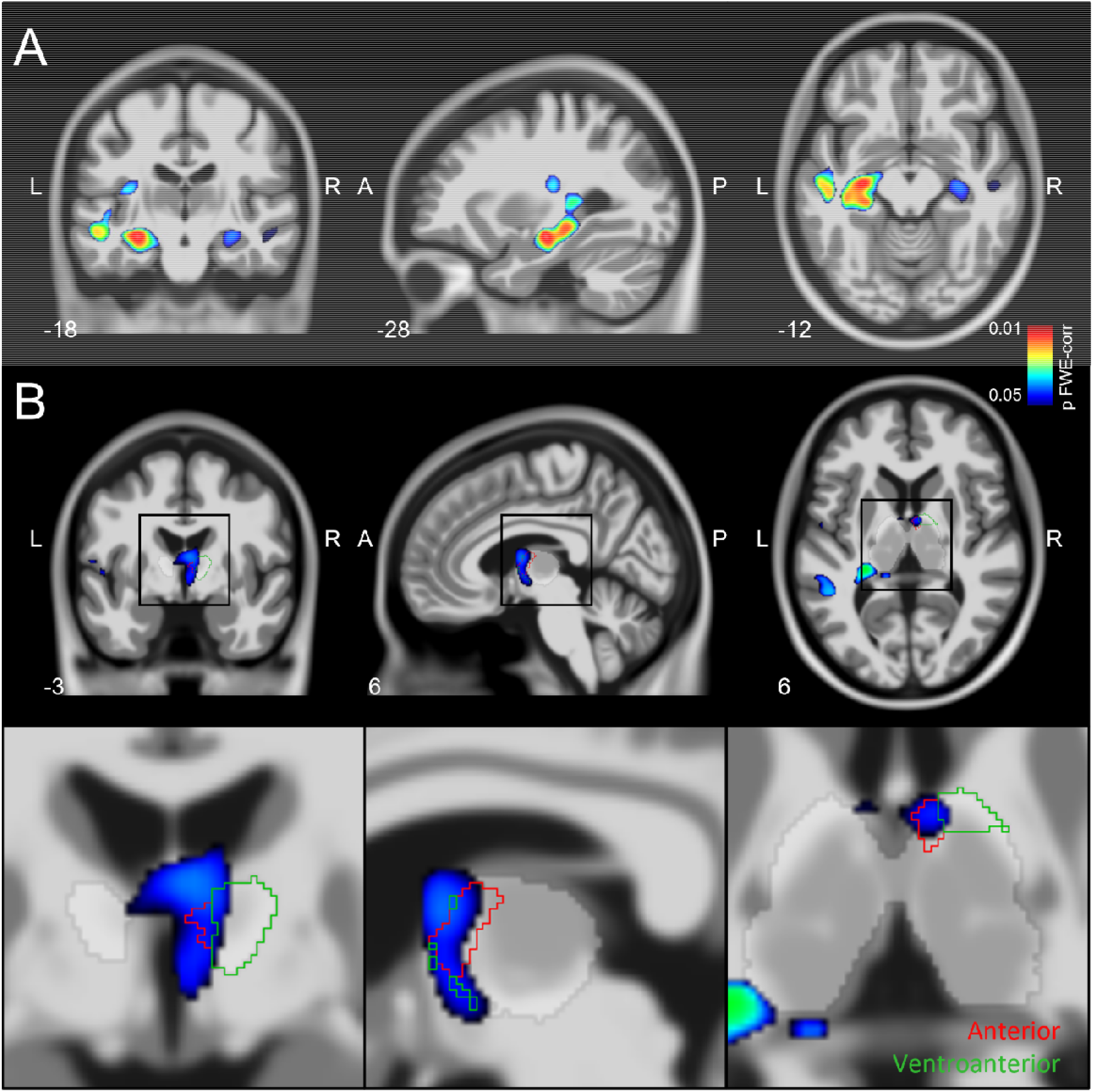
Grey matter volume correlates with episodic learning rate in older adults. The positive correlation has been overlaid on a canonical T1 image (thresholded at p < 0.05 FWE-corr) to show a significant effect in A) hippocampus bilaterally and left superior temporal gyrus and B) right anterior (red) and ventroanterior (green) (thalamic ROIs in the inset come from the AAL3 atlas (Rolls et al., 2020)). The coordinates of the sections are given in mm. L: left, R: right, A: anterior, P: posterior.

### White matter microstructure (TBSS)

We first examined FA as a marker of WM integrity. We found a bilateral network of temporal, parietal and occipital tracts showing a positive association with episodic learning rate. Among tracts showing significant positive correlations were the bilateral anterior thalamic radiation (ATR), fornix, inferior fronto-occipital fasciculus (IFOF), inferior longitudinal fasciculus (ILF), superior longitudinal fasciculus (SLF), and uncinate (Figure 3A-B, Supplementary Table 2, Supplementary Figure 2). Next, we examined MD and found a negative association between a similar network of bilateral tracts and FCSRT learning rate, including bilateral ATR, corticospinal tract, forceps major and minor, cingulum (cingulate), IFOF, ILF, SLF, uncinate and fornix (Figure 3C-D, Supplementary Figure 2, Supplementary Table 2).

**Figure 3.**
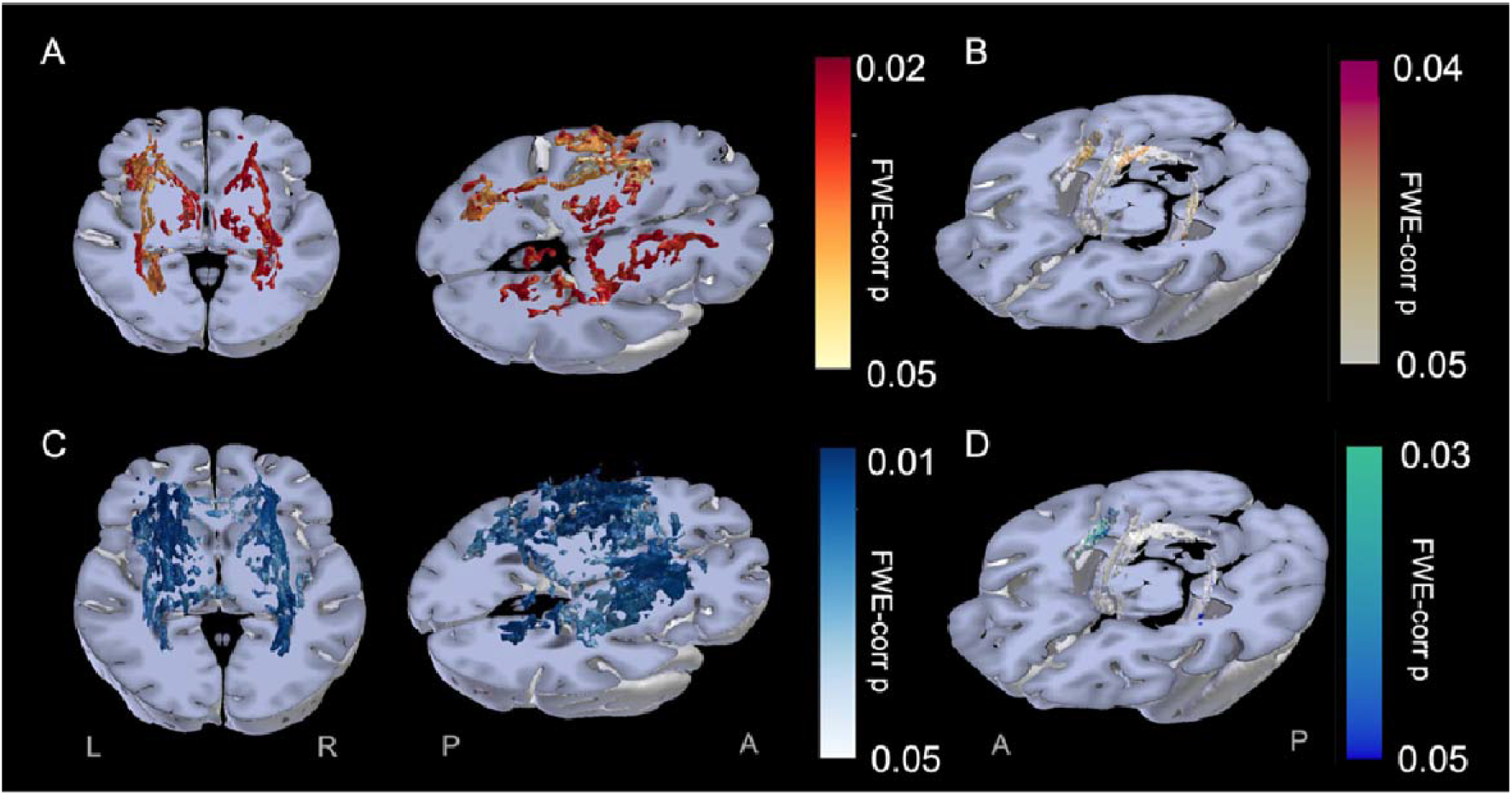
Extensive network of white matter microstructure integrity is related to episodic learning rate in older adults. A. Positive correlation between FA and learning rate (warm colours; p < 0.05 FWE-corr). B. FA effects overlaid on the fornix. C. Negative correlation between MD and learning rate (cold colours; p < 0.05 FWE-corr). D. MD effect overlaid on the fornix.

## Discussion

Our results show that women and individuals with more years of formal education had a faster episodic learning rate, that it declined with age, and that this rate was associated with neuroanatomical structural properties. We found a positive correlation between GM volume and episodic learning rate, where participants with greater volume in hippocampus, anterior thalamic nucleus and left superior temporal gyrus learned at a faster rate than those with lower volume. Furthermore, we found that FA was positively associated with episodic learning rate in an extended network including limbic tracts, indicating that the structural integrity of these tracts indexed learning ability. A complementary negative association was observed for MD, in similar tracts, such that decreased MD was associated with a faster episodic learning rate. The converging GM and WM findings suggest that structural properties of the hippocampal-anterior thalamic circuit contribute to learning ability in ageing and may potentially inform age-related decline in encoding (Friedman et al., 2007; Morcom et al., 2003).

Previous research on neural substrates of cognitive decline in ageing has shown a hippocampal volume decline with age that correlated with memory performance and with FCSRT recall specifically (Zammit et al., 2017). The presence of hippocampal volume findings with relation to both episodic learning rate and recall components of the FCSRT task suggests the hippocampus may be involved in these two separate processes, both of which are impaired in ageing. The more pronounced effect we observed in the left hippocampus is in accordance with previous VBM findings of verbal memory tasks (Ezzati et al., 2016), and the general lateralisation of verbal functions. Our results, therefore, extend previous research on the relationship between hippocampal volume and memory decline in ageing, showing episodic learning rate is also indexed by hippocampal volume. Note that it is unlikely that these effects reflect memory function in general, given that delayed free recall was included as a covariate in our model.

Structural properties of extra-hippocampal limbic regions were also associated with learning ability. Our GM thalamic findings indicate a correlation between episodic learning rate and the right anterior thalamic nucleus, extending to right ventroanterior and right ventrolateral nuclei. The anterior thalamic nuclei have been suggested to play an important role in learning and memory (Aggleton et al., 2010; Sweeney-Reed et al., 2021; Winocur, 1985), that extend beyond its established role in spatial processing (Nelson, 2021; Wolff and Vann, 2019). For example, fMRI studies in younger adults suggest that the activation of the anterior thalamic nuclei supports recognition memory performance (Pergola et al., 2013) and evidence from intracranial EEG studies indicates theta-synchronisation between anterior thalamus and frontal and parietal regions supporting successful memory formation (Sweeney-Reed et al., 2014). Furthermore, and in line with our results, Leszczyński and Staudigl (2016) posited that the anterior thalamus might modulate information flow, via attention allocation, to support learning. Taken together with the increased hippocampal volume, which was related to a better learning rate, our results indicate that the limbic system may play an important role in learning ability in ageing and might explain some of the impairments in navigating in a novel environment (Grzeschik et al., 2021), and impaired learning strategies observed in mild cognitive impairment (Ribeiro et al., 2007).

The fornix is a major hippocampal input/output pathway and has been associated with visuo-spatial learning across species (Buckley et al., 2008; Hodgetts et al., 2020; Hofstetter et al., 2013). The fornix links the hippocampus with the anterior thalamic nuclei directly and via the mammillary bodies (Aggleton et al., 2010, 1986), with both the hippocampus and the anterior thalamic nuclei showing grey matter volume relationships with episodic learning rate.

Furthermore, we found that fornix integrity, as captured by bilateral FA and MD, correlated with episodic learning rate in older adults. Together with previous findings linking fornix integrity to recall performance on the FCSRT task (Hartopp et al., 2019; Metzler-Baddeley et al., 2011), our results extend its role in memory processes, indicating that the fornix also supports verbal episodic learning. We also found that the WM integrity of the ATR was correlated with learning rate. The ATR connects the anterior and dorsomedial thalamus with the prefrontal cortex (Grodd et al., 2020), which has been suggested to play a role in learning rate (McGuire et al., 2014).

In addition to changes in limbic GM volume and WM microstructure, we found episodic learning rate was associated with broader changes within bilateral WM tracts connecting occipital-temporal-frontal regions (ILF, SLF, IFOF). This result might point toward an overall WM microstructure effect, as previously noted in age-related cognitive decline (de Mooij et al., 2018; Farokhian et al., 2017; Grieve et al., 2007; Molloy et al., 2021; Rizvi et al., 2020). With respect to specific cognitive functions, ILF and SLF have been shown to relate to memory performance in normal ageing (Sasson et al., 2013). ILF and IFOF facilitate the flow of visual information up the visual stream (Rokem et al., 2017), and IFOF has been associated with semantic processing (Duffau, 2008), potentially supporting learning performance in our task. Therefore, the observed relationship between microstructure of these tracts and learning ability might reflect a more general aspect of cognitive ability.

Finally, it is important to note some limitations of the current study; we analysed data from a cross-sectional cohort of healthy older adults. As GM and WM properties and memory function both deteriorate with age, future longitudinal studies would be needed to better understand the relationship between learning ability and structural changes as ageing progresses and eliminate age-related confounds in cross-sectional studies (Elliott, 2020). We used the learning rate across trials in an established neuropsychological memory task (FCSRT) as a measure for episodic learning; it would be interesting to examine neural correlates of learning rates in tasks such as error-driven and statistical learning (Herff et al., 2019; Nassar et al., 2016; Samanez-Larkin et al., 2012), as well as consider learning ability as a potential cognitive phenotype in pathological ageing. Finally, future research with hippocampal subfield resolution could examine their differential contribution to episodic learning rate. It would be interesting to explore whether volumetric effects are more pronounced in the subiculum, the principal source of hippocampal projections to the anterior thalamus and mammillary bodies (Hartopp et al., 2019).

In conclusion, in a cross-sectional cohort of healthy older adults, we found learning rate on the FCSRT task was positively associated with extensive GM and WM structural effects including the hippocampus, fornix and anterior thalamic nucleus, structures part of the limbic system. Furthermore, there was a positive correlation between episodic learning rate and long-range WM tracts (ILF, SLF, IFOF). Our findings indicate that episodic learning rate is associated with key anatomical structures implicated in memory function, and therefore may inform further exploration of the relationship between episodic learning rate and retrieval in ageing.

## Supplementary Materials

**Supplementary Table 1.** Coordinates of grey matter volume effects in the VBM analysis. The MNI coordinates for the global maximum and local maxima of each cluster are indicated in mm for the three sections in space (x, y and z). Neuroanatomical labels from the AAL3 atlas are indicated. P(FWE-corr): Family Wise Error corrected p-value, TFCE: Threshold-Free Cluster Enhancement local spatial support, p (unc): uncorrected p-value.

| Number of<br>voxels in<br>cluster | p (FWE-corr) | TFCE | p (unc) | x | y | z | AAL3 labels |
| --- | --- | --- | --- | --- | --- | --- | --- |
| 2287 | 0.007 | 1337.9 | 0 | -28 | -18 | -12 | Hippocampus_L<br>Extended to the left<br>superior temporal gyrus |
|  | 0.008 | 1318.49 | 0 | -40 | -30 | -8 | Hippocampus_L |
|  | 0.008 | 1308.5 | 0 | -26 | -31 | -8 | Hippocampus_L |
| 184 | 0.032 | 999.29 | 0.002 | 4 | -3 | 12 | Thal_VL_R |
|  | 0.038 | 959.07 | 0.001 | 4 | -5 | -2 | Thal_VA_R |
| 168 | 0.033 | 994.55 | 0.001 | 30 | -19 | -12 | Hippocampus_R |
|  | 0.045 | 922.07 | 0.002 | 20 | -23 | -16 | Parahippocampal_R |
| 46 | 0.044 | 928.22 | 0.001 | 20 | -67 | 22 | Cuneus_R |
|  | 0.049 | 904.13 | 0.001 | 18 | -59 | 16 | Calcarine_R |
| 64 | 0.046 | 915.85 | 0.002 | 52 | -19 | -12 | Temporal_Mid_R |
|  | 0.047 | 911.21 | 0.002 | 44 | -11 | -18 | Hippocampus_R |
| 22 | 0.048 | 909.89 | 0.002 | -24 | 18 | -18 | OFCpost_L |
| 7 | 0.049 | 903.11 | 0.001 | 14 | -28 | -24 | Cerebellum_3_R |

**Supplementary Table 2.** Brain clusters for correlation between episodic learning rate and FA and MD values in the main analysis. P(FWE-cor): Family Wise Error corrected p-value. Neuroanatomical labels from the JHU white matter atlas are indicated.

| Number of voxels in cluster | p (FWE-corr) | Main tracts |
| --- | --- | --- |
| Positive correlation with FA values |  |  |
| 4917 | 0.025 | IFOF, SLF, fornix, ATR_R |
| 3123 | 0.036 | IFOF, ILF, SLF, fornix, ATR_L |
| 1310 | 0.036 | IFOF, ILF, forceps major_R |
| 359 | 0.047 | ILF, forceps major_L |
| 239 | 0.044 | SLF_L |
| 128 | 0.047 | Corpus callosum |
| 61 | 0.049 | Forceps minor |
| 8 | 0.05 | Corpus callosum |
| Negative correlation with MD values |  |  |
| 9921 | 0.013 | IFOF, ILF, SLF, ATR_R, forceps minor, corpus callosum |
| 7467 | 0.015 | IFOF, ILF, SLF, fornix, ATR_L, corpus callosum |
| 3 | 0.05 | Corticospinal tract |
| 1 | 0.05 |  |

**Supplementary Figure 1.**
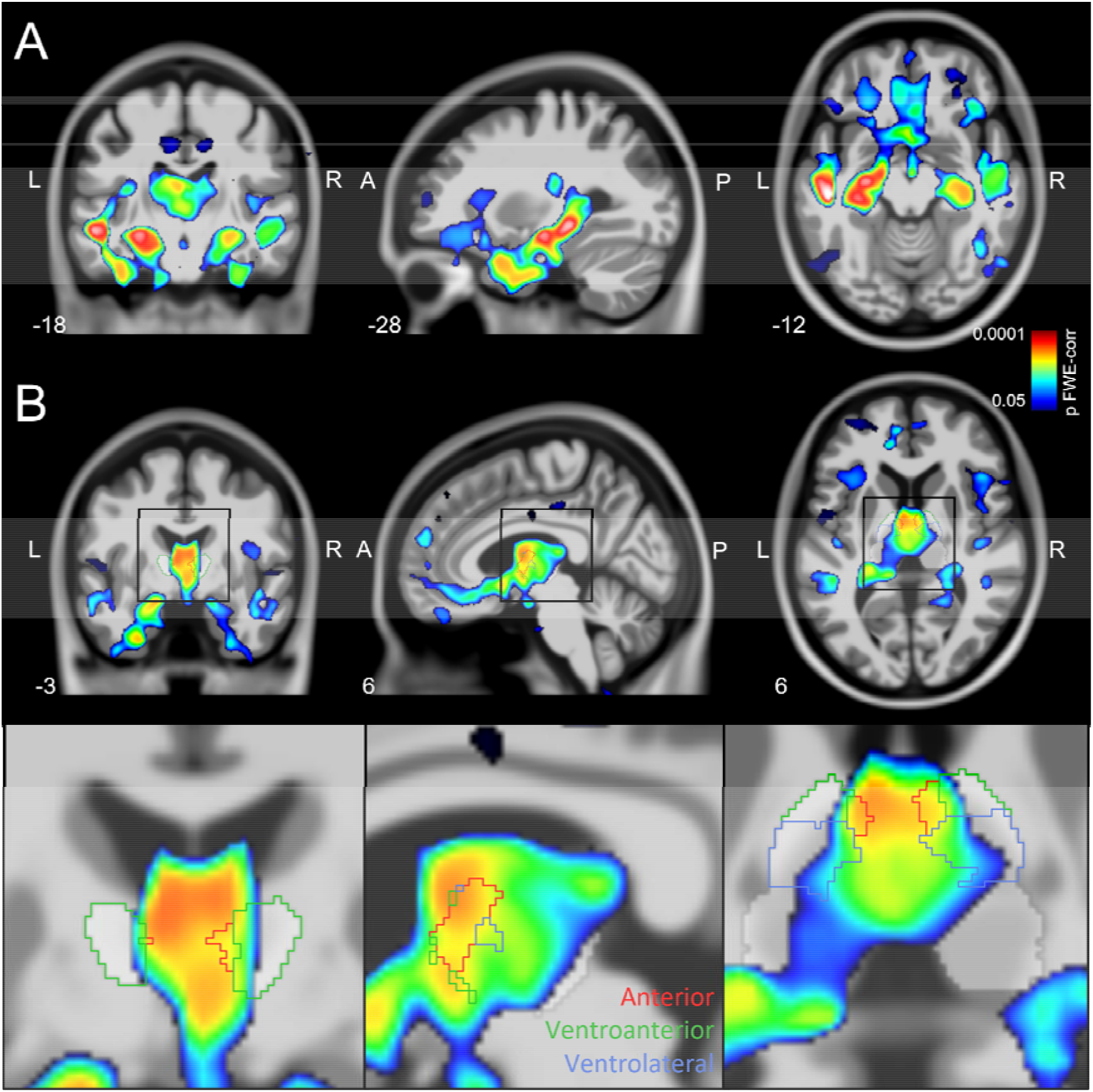
Grey matter volume correlates with learning rate in older adults with an alternative statistical model. For this model, age and education were introduced in the model whereas the delayed FCSRT score was removed. The positive correlation has been overlaid on a canonical T1 image (thresholded at p < 0.05 FWE-corr) to show a significant effect in A hippocampus bilaterally and B thalamus, right anterior (red), ventroanterior (green) and ventrolateral thalamic nuclei (blue) (thalamic ROIs in the inset come from the AAL3 atlas (Rolls et al., 2020). The coordinates of the sections are given in mm. L: left, R: right, A: anterior, P: posterior.

**Supplementary Figure 2.**
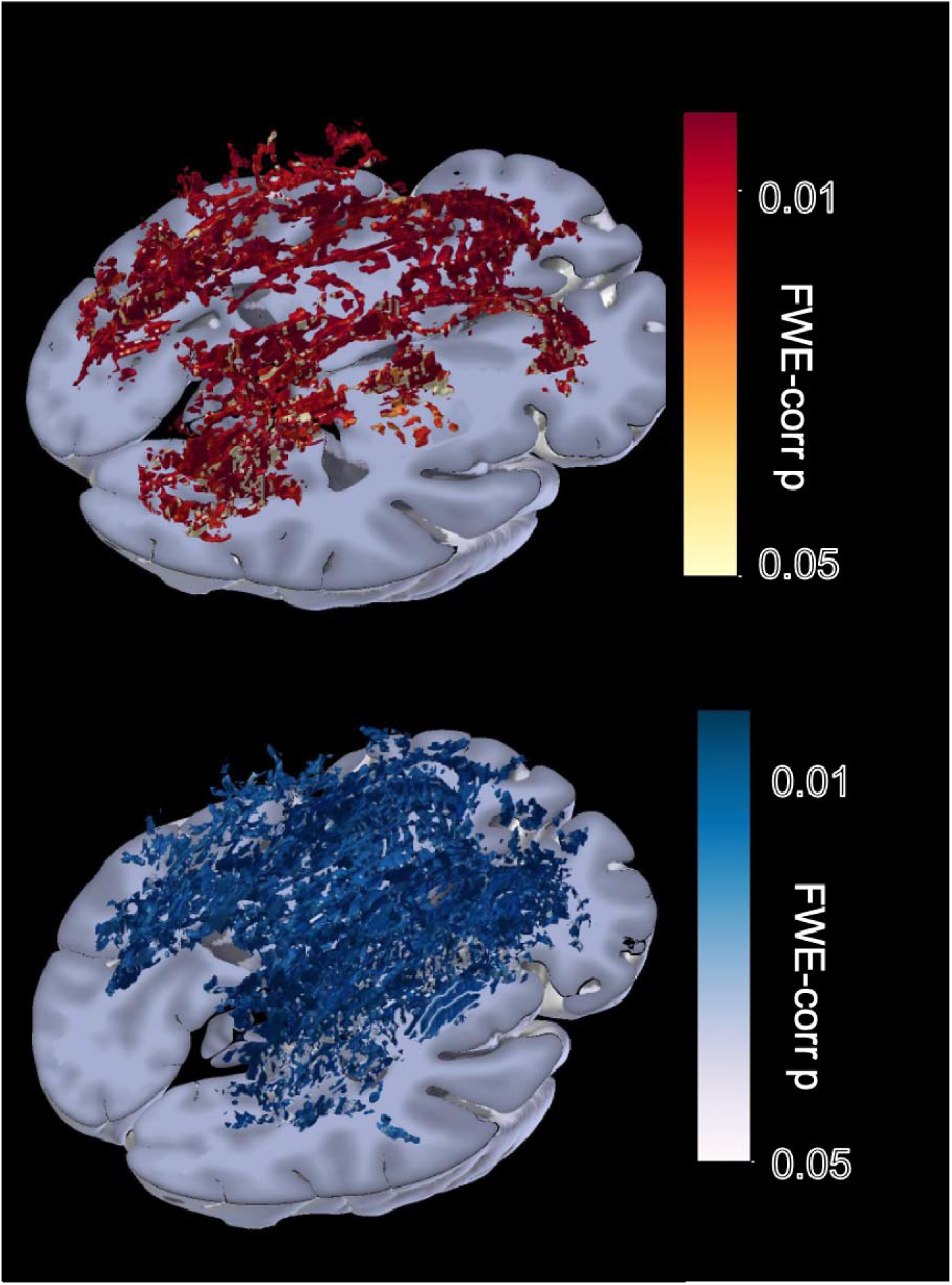
Extensive network of white matter microstructure integrity is related to learning rate in older adults with an alternative statistical model. For this model, age and education were introduced in the model whereas the delayed FCSRT score was removed. A. Positive correlation between FA and learning rate (warm colours; p < 0.05 FWE-corr). B. FA effects overlaid on the fornix. C. Negative correlation between MD and learning rate (cold colours; p < 0.05 FWE-corr). D. MD effect overlaid on the fornix.

